# Visible blue light inactivates SARS-CoV-2 variants and inhibits Delta replication in differentiated human airway epithelia

**DOI:** 10.1101/2022.01.25.477616

**Authors:** Jacob Kocher, Leslee Arwood, Rachel C. Roberts, Ibrahim Henson, Abigail Annas, David Emerson, Nathan Stasko, M. Leslie Fulcher, Marisa Brotton, Scott H. Randell, Adam S. Cockrell

## Abstract

The emergence of SARS-CoV-2 variants that evade host immune responses has prolonged the COVID-19 pandemic. Thus, the development of an efficacious, variant-agnostic therapeutic for the treatment of early SARS-CoV-2 infection would help reduce global health and economic burdens. Visible light therapy has the potential to fill these gaps. In this study, visible blue light centered around 425 nm efficiently inactivated SARS-CoV-2 variants in cell-free suspensions and in a translationally relevant well-differentiated tissue model of the human large airway. Specifically, 425 nm light inactivated cell-free SARS-CoV-2 variants Alpha, Beta, Delta, Gamma, Lambda, and Omicron by up to 99.99% in a dose-dependent manner, while the monoclonal antibody bamlanivimab did not neutralize the Beta, Delta, and Gamma variants. Further, we observed that 425 nm light reduced virus binding to host ACE-2 receptor and limited viral entry to host cells *in vitro*. Further, the twice daily administration of 32 J/cm^2^ of 425 nm light for three days reduced infectious SARS-CoV-2 Beta and Delta variants by >99.99% in human airway models when dosing began during the early stages of infection. In more established infections, logarithmic reductions of infectious Beta and Delta titers were observed using the same dosing regimen. Finally, we demonstrated that the 425 nm dosing regimen was well-tolerated by the large airway tissue model. Our results indicate that blue light therapy has the potential to lead to a well-tolerated and variant-agnostic countermeasure against COVID-19.

## Introduction

In late 2019, the severe acute respiratory syndrome coronavirus 2 (SARS-CoV-2), the causative agent of Coronavirus Disease 2019 (COVID-19), emerged in Wuhan, China and rapidly spread around the globe, resulting in nearly 237 million confirmed cases and five million deaths (Ritchie et al., 2020). Due to hot spots of uncontrolled spread, novel variants have emerged displaying various combinations of increased replication, increased virulence, increased transmission, and the ability to evade immune response from previous infections or vaccination (Harvey et al., 2021; Krause et al., 2021). For example, the Beta variant was notable for its ability to evade monoclonal antibodies and serum neutralizing antibody responses (Wang et al., 2021). Similarly, the SARS-CoV-2 Delta and Omicron variants demonstrated combinations of increased transmission, virulence, and immune evasion and rapidly became the dominant global strains in early and late 2021, respectively, unleashing fresh “waves” of infections, hospitalizations, mortality, and economic instability (Y. Liu et al., 2021; Planas et al., 2021; Pouwels et al., 2021; Ren et al., 2022; Xu et al., 2021). Rising COVID-19 cases in regions across the world provides ample opportunity for new variants (e.g. Lambda and Mu) to emerge and further threaten global recovery (H. Liu et al., 2021; Ritchie et al., 2020).

Although much effort has been placed on the development of vaccines and therapeutics to reduce the prevalence and disease severity of COVID-19, additional countermeasures to return the world to pre-pandemic conditions are still needed. Several efficacious vaccines derived from the original, parental SARS-CoV-2 strain have been distributed, but <50% of the worldwide population is fully vaccinated (Ritchie et al., 2020), and recent studies have suggested waning protection in vaccinated individuals (Levin et al., 2021; Naaber et al., 2021). Therapeutic development, while also encouraging, has lagged. Several approaches have shown promise in laboratory or small-scale clinical studies with the best success in early onset disease and via a combination of therapeutic modalities (Caly et al., 2020; Liu et al., 2020; McCullough et al., 2021). The only therapeutics currently authorized by the Food and Drug Administration are therapeutic monoclonal antibodies, molnupiravir, nirmatrelvir/ritonavir, and remdesivir, which have limitations including intravenous infusions, cost, susceptibility to variant escape, and questionable utility outside of specialized care facilities (Beigel et al., 2020; Jayk Bernal et al., 2021; Kumar et al., 2021). These limitations highlight a critical need for additional therapeutics in the fight against COVID-19, particularly for a variant-agnostic countermeasure that can be easily administered in the home setting. Visible light has the potential to fulfill this glaring need. McNeil and coworkers previously demonstrated the utility of visible 425 nm light targeted to the oropharynx and surrounding tissue as a possible at-home treatment for mild-to-moderate COVID-19 (Stasko et al., 2021a), but the broad utility of targeted blue light against SARS-CoV-2 variants of concern has not been established.

We previously reported that 425 nm blue light inactivates SARS-CoV-2 WA1 in cell-free and cell-associated formats at doses that are well-tolerated in a human tracheobronchial tissue model (Stasko et al., 2021b) and that also induce host IL-1α and IL-1β gene expression in a human oral epithelial tissue model (Stasko et al., 2021a). Additionally, we demonstrated that human airway models can tolerate up to 256 J/cm^2^ of 425 nm light given in a twice daily 32 J/cm^2^ regimen for four days (Stasko et al., 2021b). The present study builds upon those seminal findings; herein we show that 425 nm blue light is capable of consistently inactivating each of the major SARS-CoV-2 variants tested to date. Further, we show that twice daily repeat dosing inhibits the replication of the Beta and Delta SARS-CoV-2 variants in a translationally relevant three-dimensional, differentiated model of human tracheobronchial epithelia. Our findings reveal the utility of safe, visible light as a variant-agnostic therapeutic for acute SARS-CoV-2 infection and detail its potential in treating infectious caused by SARS-CoV-2 variants.

## Methods

### Cells, tissues, and viruses

Vero E6 and A549 cells were purchased from ATCC. Vero E6 cells were maintained as previously described (Stasko et al., 2021b). A549 cells were maintained according to ATCC instructions. Human adenovirus type 5 (VR-5) was acquired from the ATCC and passaged as previously described (Stasko et al., 2021b).

SARS-CoV-2 isolates WA1 (NR-52281), Alpha (NR-54000), Beta (NR-54009), Delta (NR-55611), Gamma (NR-54982), Lambda (NR-55654), and Omicron (NR-56461) were obtained through BEI Resources, National Institute of Allergy and Infectious Diseases, National Institutes of Health and passaged as previously described (Stasko et al., 2021b). All work with replication-competent SARS-CoV-2 was conducted with adherence to established safety guidelines at the CDC-certified biosafety level-3 facility at EmitBio, Inc.

The MatTek EpiAirway™ large airway epithelial models (donors AIR-100 and TBE-14) were acquired and handled as previously described (Stasko et al., 2021b). Primary human tissue models derived from large airway epithelial cells were also acquired from the Marsico Lung Institute Tissue Procurement and Cell Culture Core facility at the University of North Carolina at Chapel Hill (Donor DD065Q). The cells were cultured and differentiated on Millicell CM™ inserts in UNC air-liquid interface media as previously described (Fulcher and Randell, 2013). Cells were delivered in 12-well plates with agarose embedded in the basal compartment and revived as previously described (Stasko et al., 2021b). Cells were allowed to recover for 3-5 days prior to experimental initiation. Antiviral assays were conducted as previously described (Stasko et al., 2021b).

### Plaque assay

Infectious viral titers for SARS-CoV-2 were enumerated via plaque assay as previously described (Stasko et al., 2021b). SARS-CoV-2 WA1 plaque assays were fixed and stained at 4 days post-infection and SARS-CoV-2 variants were fixed and stained at 5 days post-infection. Human adenovirus plaque assays were conducted on A549 cells with 1.2% colloidal microcrystalline cellulose overlay and developed 5 days post-infection.

### 425 nm light plaque reduction neutralization test (PRNT)

Virus stocks were diluted to 2×10^5^ PFU/mL (variants), 2×10^6^ PFU/ml (WA1), or 2×10^7^ PFU/mL (adenovirus) in media and illuminated with the indicated doses of 425 nm light as previously described (Stasko et al., 2021b). Illuminated viral suspensions were titered via plaque assay as above.

### Monoclonal antibody PRNT

Bamlanivimab (mAb LY-CoV555) was diluted to 4 μg/mL and serially diluted 1:2. SARS-CoV-2 stocks were diluted to 250 PFU/50 μl and 60 μL was added to each antibody dilution (final concentrations 2 μg/ml to 0.00391 μg/mL). Virus-antibody combinations were incubated for 1 hour at 37°C/5% CO_2_ and then titered via plaque assay as above. Plaques were counted for each antibody dilution and for virus input; 50% and 90% neutralization cut off was determined based on the virus input.

### Binding ELISA

Fifty μL (0.25 μg/mL) of human ACE-2 was added to Nickel Coated Assay Plates and incubated at room temperature for 1 h. SARS-CoV-2 Beta was diluted to 2×10^5^ PFU/mL and illuminated (0 J/cm^2^, 15 J/cm^2^, and 90 J/cm^2^) as described above and then diluted 1:10000 in PBS. ACE-2 was removed from the wells and 50 μL of each virus suspension was added to each well and incubated for 1 h at room temperature. Fifty μL of 1:3000 diluted mouse anti-SARS-CoV-2 Spike Protein S1 monoclonal antibody (GT623, Invitrogen) in 1% BSA-PBS blocking buffer was added and incubated at room temperature for 30 min. Fifty μL of HRP-conjugated goat anti-mouse IgG diluted 1:2000 in 1% BSA-PBS blocking buffer was added to each well and incubated for 30 minutes at room temperature. Plates were washed 3× with 0.5% PBS-Tween20 between each step above. Plates were developed with 1-Step Ultra TMB-ELISA for 15-30 minutes. Reactions were stopped with 2M sulfuric acid and read at 450 nm.

### Cell entry qPCR

SARS-CoV-2 Beta was diluted to 2×10^5^ PFU/mL and illuminated with 425 nm light (0 J/cm^2^, 15 J/cm^2^, and 90 J/cm^2^) as above. Vero E6 cells were inoculated with 200 μL of illuminated virus suspension and incubated for 1 h at 37°C/5% CO_2_. The inoculum was removed and cells washed 2× with PBS and 500 μL culture media (high glucose DMEM + 5% FBS (Gibco) + 1% antibiotic-antimycotic) was added to each well. Cells were incubated at 37°C/5% CO_2_ and washed 2× with PBS prior to total RNA extraction. Total RNA was extracted at 3 hpi and 24 hpi with the RNeasy Mini Total RNA extraction kit (Qiagen). SARS-CoV-2 RNA was detected with the CDC assay kit (IDT) and the TaqMan Fast Virus 1-Step Master Mix (ThermoFisher) on the QuantStudio3 System (Invitrogen).

### Statistical analysis

Statistical significance for viral titers and cell cytotoxicity were determined using the Mann-Whitney ranked sum test using GraphPad Prism 8. Statistical significance is indicated by * (p ≤ 0.05) and ** (p ≤ 0.01). For statistical purposes, all samples that were under the limit of detection were set to half the assay limit of detection.

## Results

### Inactivation of SARS-CoV-2 variants assessed via plaque reduction neutralization tests (PRNT)

We previously demonstrated that 425 nm light inactivated highly pathogenic betacoronaviruses in cell-free suspensions (Stasko et al., 2021b). In this study, we expanded upon these findings by evaluating whether the same doses of 425 nm light inactivated a panel SARS-CoV-2 variants (Alpha, Beta, Delta, Gamma, Lambda, and Omicron) using the plaque reduction neutralization test (PRNT) (Figure 1). Similar to our previous report (Stasko et al., 2021b), 425 nm light reduced SARS-CoV-2 infectious titers in a dose-dependent manner: >1 log_10_ at 7.5 J/cm^2^, >2 log_10_ at 15 J/cm^2^, >3 log10 at 30 J/cm^2^, and >4 log_10_ at 60 J/cm^2^ (Figures 2A-2G). We observed no inactivation of the non-enveloped DNA virus human adenovirus 5 (Figure 2H); we previously demonstrated that similar findings with human rhinovirus (Stasko et al., 2021b), suggesting these light doses do not damage viral RNA or viral DNA and work in an envelope-dependent mechanism. Additionally, we observed that basal media composition and sera supplementation had no impact on inactivation kinetics (Supplementary Figure 1), suggesting direct inactivation of the SARS-CoV-2 virion independent of potential differences in media composition (e.g. increased amino acids or lot-specific serum factors).

**Figure 1.**
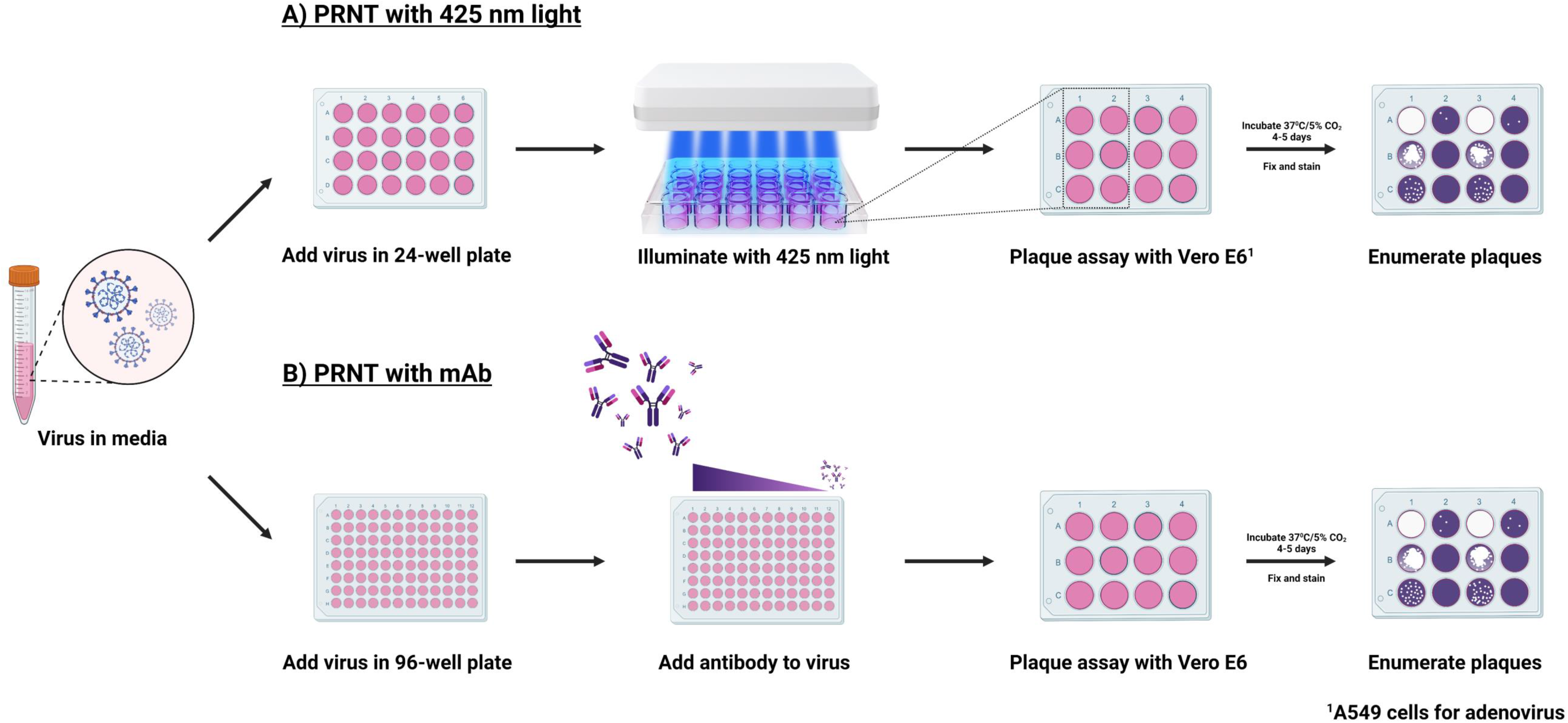
Scheme of Plaque Reduction Neutralization Test (PRNT) for 425 nm light and bamlanivimab. Biological light units were configured to evenly distribute light over the entire surface area of a 24-well plate and used to evaluate various energy densities in the classic PRNT assay method to measure inactivation of cell-free SARS-CoV-2. SARS-CoV-2 variants were diluted and illuminated with (A) a fixed dose of 425 nm light per experiment or (B) incubated with serially diluted bamlanivimab prior to infectious titer enumeration with plaque assay.

**Figure 2.**
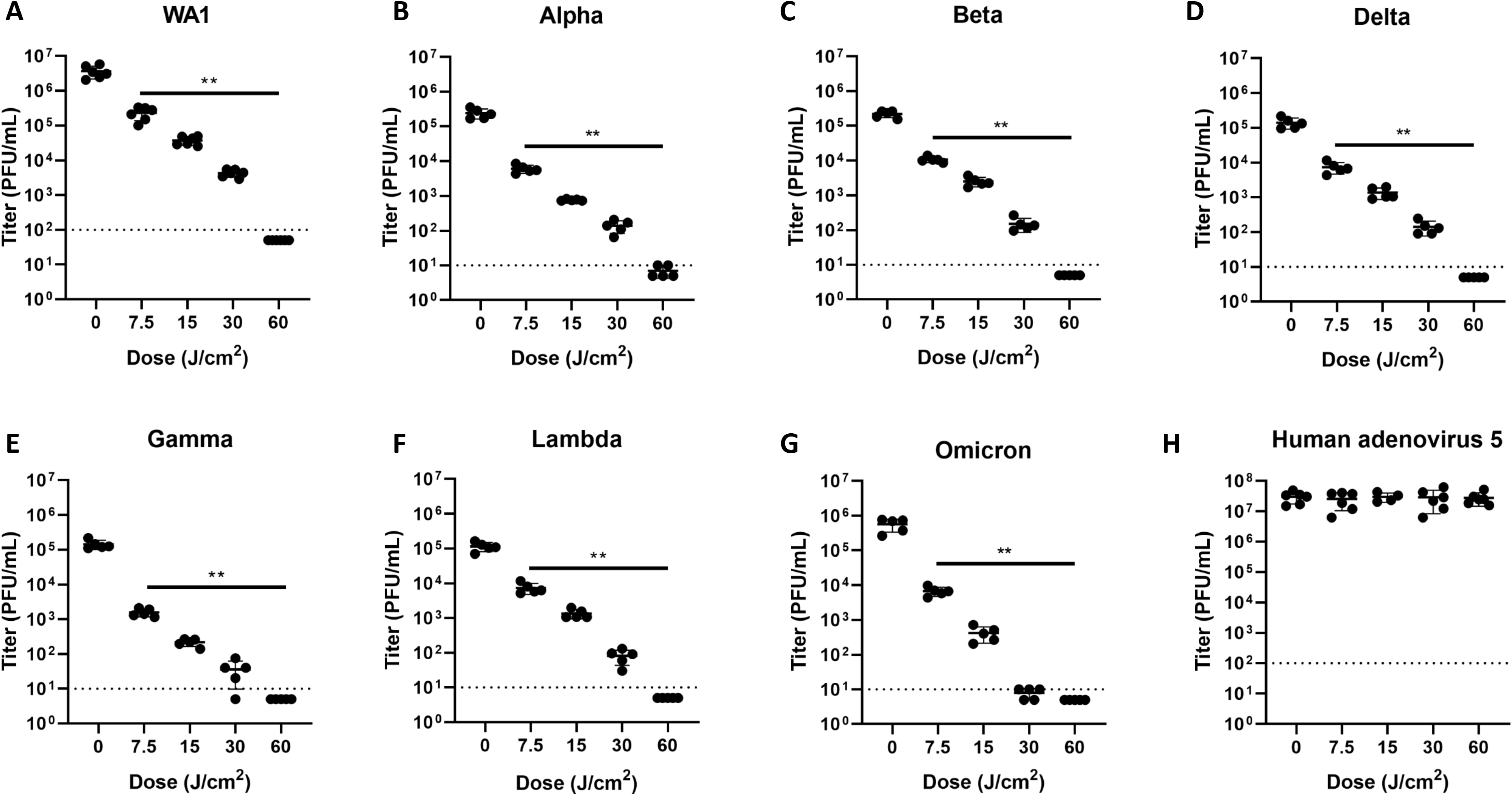
425 nm light inactivates up to 99.99% of cell-free SARS-CoV-2 variants in a dose-dependent manner, but not human rhinovirus or human adenovirus. Cell-free suspensions of SARS-CoV-2 (A) WA1, (B) Alpha, (C) Beta, (D) Delta, (E) Gamma, (F) Lambda, and (G) Omicron and (H) human adenovirus type 5 were illuminated with 425 nm light and enumerated via plaque assay. Data presented are mean viral titers +/− SD (n = 5-6). Statistical significance was determined via Mann-Whitney ranked sum test and is indicated by * (p≤0.05) and ** (p≤0.01).

Previous studies have demonstrated that several variants evade neutralization by convalescent plasma and monoclonal antibody therapies (e.g. bamlanivimab) (Cao et al., 2021; Cele et al., 2021; H. Liu et al., 2021; Planas et al., 2021; VanBlargan et al., 2022; Widera et al., 2021). Thus, bamlanivimab was included as a comparison to illustrate the evasion ability of replication-competent SARS-CoV-2 variants in our laboratory. In the present study, we observed that Beta, Delta, and Gamma evaded neutralization by bamlanivimab, but WA1 and Alpha did not, which is consistent with previous reports (Planas et al., 2021; Widera et al., 2021). When comparing the PRNT_50_ and PRNT_90_ titers for 425 nm light and bamlanivimab (Table 1), we observed that the PRNT_50_ and PRNT_90_ bamlanivimab titers for Beta, Delta, and Gamma were ≥ 128 fold greater than WA1. Conversely the PRNT_50_ and PRNT_90_ doses of blue light for WA1, Alpha, Beta, Delta, Gamma, Lambda, and Omicron variants had similar PRNT_50_ and PRNT_90_ values, indicating that SARS-CoV-2 variants remain susceptible to 425 nm light inactivation.

**Table 1.** Bamlanivimab and 425nm light PRNT_50_ and PRNT_90_ values.

|  | Bamlanivimab |  |  |  | 425nm light |  |  |  |
| --- | --- | --- | --- | --- | --- | --- | --- | --- |
|  | PRNT<sub>50</sub> |  | PRNT<sub>90</sub> |  | PRNT<sub>50</sub> |  | PRNT<sub>90</sub> |  |
|  | [µg/mL] | Fold<br>change<br>over WA1 | [µg/mL] | Fold<br>change<br>over WA1 | J/cm<sup>2</sup> | Fold<br>change<br>over WA1 | J/cm<sup>2</sup> | Fold<br>change<br>over WA1 |
| **WA1** | 0.022 | 0 | 0.063 | 0 | 2.8 | 0 | 6.2 | 0 |
| **Alpha** | 0.011 | <1 | 0.045 | <1 | 2.4 | <0 | 4.7 | <0 |
| **Beta** | >2 | >256<sup>A</sup> | >2 | >256 | 1.9 | <0 | 5.3 | <0 |
| **Delta** | >1 | >128 | >2 | >256 | 2.2 | <0 | 5.6 | <0 |
| **Gamma** | >2 | >256 | >2 | >256 | 1.2 | <0 | 2.9 | <0 |
| **Lambda** | ND* | ND | ND | ND | 2.5 | <0 | 6.2 | <0 |
| **Omicron** | ND | ND | ND | ND | 2.2 | <0 | 4.0 | <0 |

\*ND = not determined, Lambda and Omicron evasion of bamlanivimab neutralization has been reported (Cao et al., 2021; H. Liu et al., 2021; VanBlargan et al., 2022)
<sup>A</sup> For samples >2 µg/mL, values were set to 4 for calculation of fold change

### 425 nm light inactivates SARS-CoV-2 through inhibition of ACE-2 binding and cellular entry

To investigate the mechanism of 425 nm light inactivation of cell-free SARS-CoV-2, cell-free SARS-CoV-2 Beta was illuminated with two doses of 425 nm light and its ability to bind ACE-2 and enter host cells was assessed (Figure 4). We selected a non-virucidal dose (15 J/cm^2^) and a high dose 50% above the reported virucidal dose (90 J/cm^2^) to ensure complete inactivation for these assays. To determine if illuminated virus maintained ACE-2 binding integrity, we conducted a human ACE-2 receptor-ligand binding assay (Figure 4A). Illumination with 425 nm light reduced SARS-CoV-2 Beta binding to ACE-2 in a dose-dependent manner, as 15 J/cm^2^ reduced binding by ~80% and 90 J/cm^2^ eliminated all ACE-2 binding.

**Figure 3.**
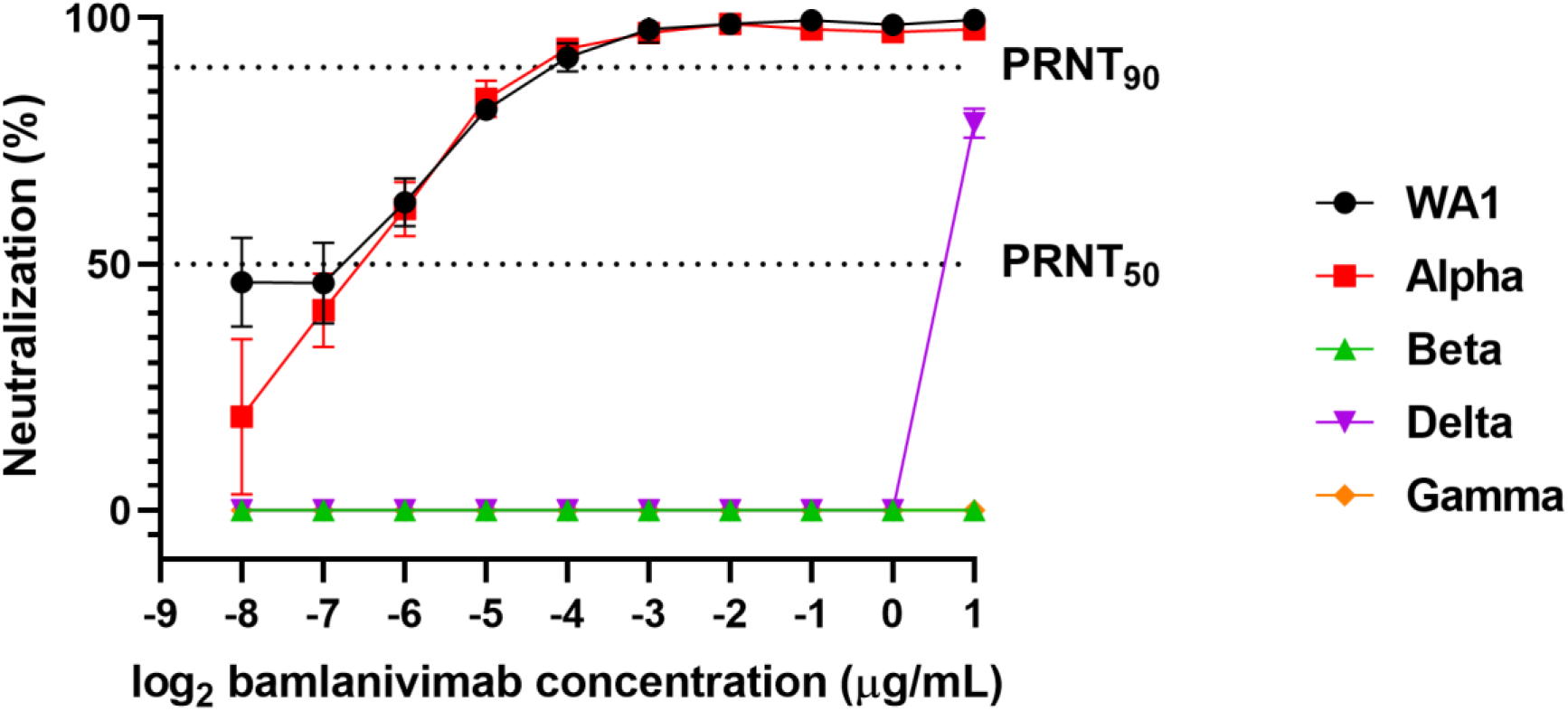
SARS-CoV-2 variants Beta, Delta, and Gamma evade neutralization by the monoclonal antibody therapeutic bamlanivimab. The neutralization capability of bamlanivimab with cell-free SARS-CoV-2 variants (WA1, Alpha, Beta, Delta, and Gamma) was determined by the plaque reduction neutralization test. Cell-free SARS-CoV-2 variants were incubated with varying concentrations of bamlanivimab and viral titers were enumerated via plaque assay. Data presented are the mean percent neutralization +/− SD (n = 4) across bamlanivimab concentrations (μg/mL).

**Figure 4.**
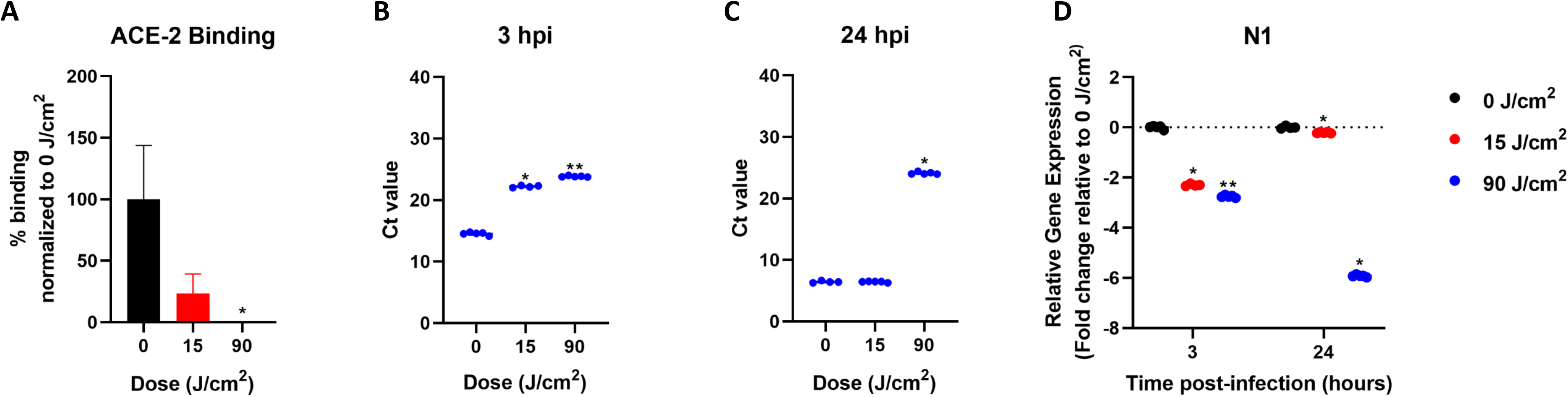
425 nm light inhibits cell-free SARS-CoV-2 cell entry in an ACE-2-dependent manner. (A) Cell-free suspensions of SARS-CoV-2 Beta were illuminated with 0 J/cm^2^, 15 J/cm^2^, or 90 J/cm^2^ of 425 nm light. Illuminated samples were evaluated for binding to ACE-2 via ELISA (n = 4). Data presented are normalized ACE-2 binding (relative to 0 J/cm^2^) +/− SEM. (B) Vero E6 cells were inoculated with cell-free suspensions of SARS-CoV-2 Beta following illumination with 0 J/cm^2^, 15 J/cm^2^, and 90 J/cm^2^ of 425 nm light. At 3 hpi and 24 hpi, total RNA was extracted from inoculated cultures for qRT-PCR analysis using the CDC N1 primer/probe set (n = 4). Data presented are mean Ct +/− SD (n = 4) for the N1 probe at (B) 3 hpi and (C) 24 hpi and (D) normalized N1 relative to host RNase P. Statistical significance was determined via the Mann-Whitney ranked sum test and is indicated by * (p≤0.05) and ** (p≤0.01).

Using the same light doses, we inoculated Vero E6 cells with virus that had been previously exposed to blue light and evaluated cell-associated SARS-CoV-2 viral replication via N1 qRT-PCR at 3 hpi (Figure 4B) and 24 hpi (Figure 4C). At 3 hpi, both doses significantly reduced detectable viral RNA compared to the unilluminated control. However, at 24 hpi, viral RNA from 15 J/cm^2^ had similar amounts of viral RNA as cells inoculated with unilluminated virus suspensions, suggesting that, while viral binding is reduced, these virions are still capable of undergoing replication. Conversely, SARS-CoV-2 illuminated with 90 J/cm^2^ of 425 nm light had significantly lower amounts of detectable viral RNA and did not change significantly from 3 hpi to 24 hpi, indicating impaired viral entry into the host cell following complete inactivation. Gene expression normalized to host RNaseP confirmed these results; 15 J/cm^2^ reduced detectable RNA by 2 logs at 3 hpi and 90 J/cm^2^ reduced detectable RNA by 2 logs and 6 logs at 3 hpi and 24 hpi, respectively (Figure 4D). We observed similar trends with the N2 qRT-PCR (Supplementary Figure 2A-C) and, as expected, did not observe a dose-dependent effect in RNaseP from Vero E6 cells (Supplementary Figure 2D-2E). These results indicate that 425 nm light inactivates SARS-CoV-2 by inhibiting viral binding and entry to the host cell.

### 425 nm light inhibits replication of the SARS-CoV-2 Beta variant in a human tissue infection model

To determine the best tissue model for screening 425 nm blue light against SARS-CoV-2-infected well-differentiated human airway epithelial tissues, we evaluated WA1 replication kinetics in multiple donor models from different suppliers (Supplementary Figure 3). SARS-CoV-2 replicated more consistently and to higher peak titers in the DD065Q model (UNC MLI) than in the AIR-100 and TBE-14 EpiAirway™ models. Additionally, the inhibition of WA1 replication following exposure to 425 nm light was evaluated in two separate model systems (DD065Q and TBE-14). In the TBE-14 model, 32 J/cm^2^ reduced WA1 titers below the limit of detection compared to the 2 log_10_ reduction observed in the DD065Q model. Since SARS-CoV-2 variants have demonstrated increased replication *in vitro* and *in vivo* (Arora et al., 2021; Cheng et al., 2021; Plante et al., 2021; Touret et al., 2021b), potentially narrowing the therapeutic window of new countermeasures, we selected the more stringent DD065Q model for the studies described herein. Thus, we treated SARS-CoV-2 Beta-infected tissues (MOI 0.1) with 32 J/cm^2^ of 425 nm light once daily (QD) or twice daily (BID) starting at 3 hpi for three days (Figure 5A). While the QD regimen reduced titers by >1 log_10_ at 72 hpi, the BID regimen reduced titers >4 log_10_ at 72 hpi (Figure 5B). Importantly, the SARS-CoV-2 titers in BID-treated tissues decreased from 24 hpi to 72 hpi, indicating the inhibition of the SARS-CoV-2 Beta replication. Additionally, we observed no light-induced cytotoxicity in time-matched, uninfected tissues after 3 days of repeat dosing (Figure 5C). These results demonstrate that a BID, but not QD, dosing regimen with 32 J/cm^2^ of 425 nm light is sufficient to inhibit SARS-CoV-2 Beta in a well-differentiated airway tissue model.

**Figure 5.**
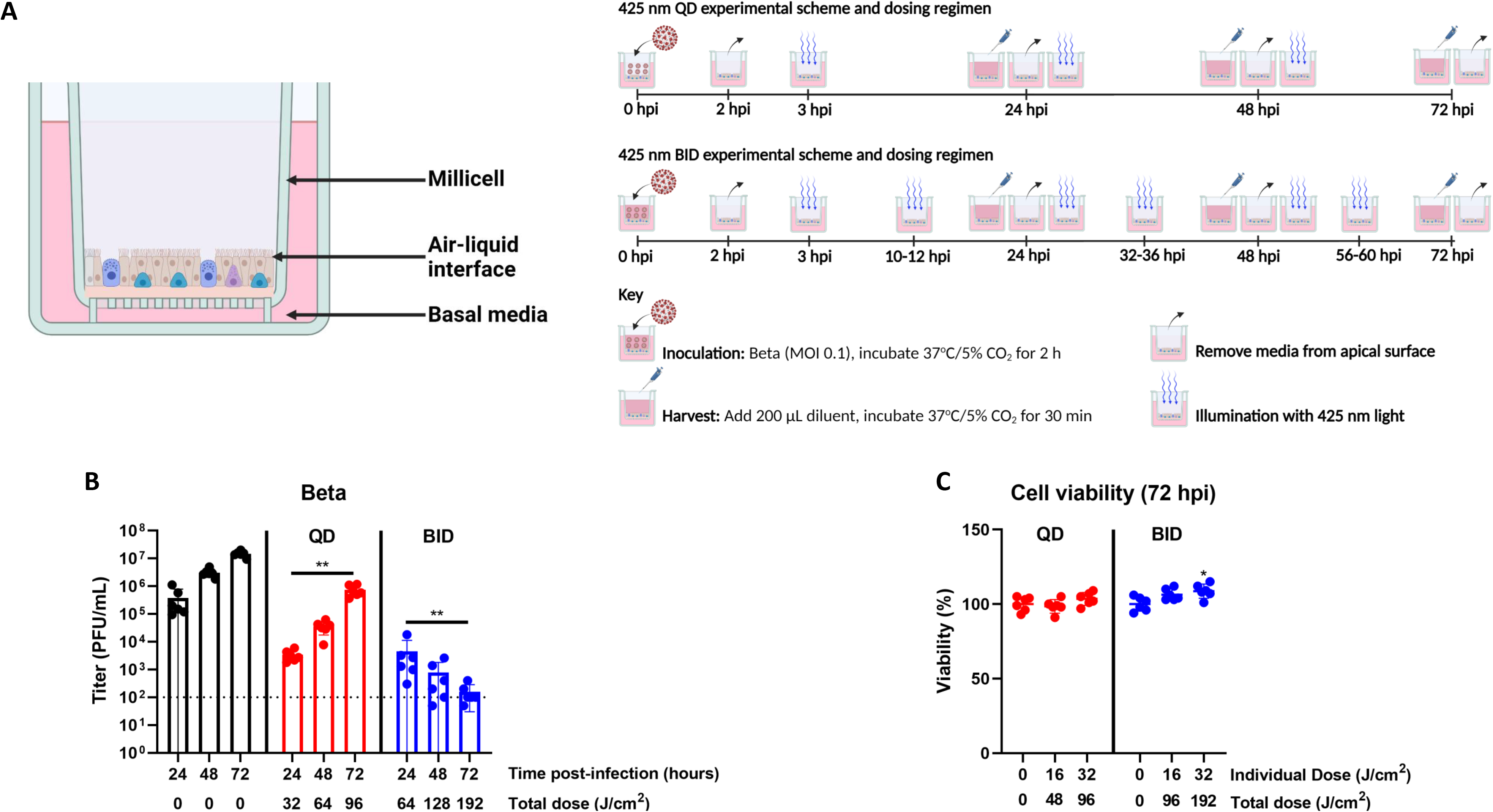
Twice daily exposures of 32 J/cm^2^ of 425 nm light inhibit productive SARS-CoV-2 Beta infection and is well-tolerated in a primary human airway tissue model. (A) Well-differentiated primary human airway tissue models were infected with SARS-CoV-2 Beta (MOI 0.1) and illuminated once (QD) or twice daily (BID) for 3 days with 32 J/cm^2^ of 425 nm light starting at 3 hours post-infection. (B) Apical rinses were collected daily and enumerated via plaque assay. Data presented are mean viral titer (PFU/mL) +/− SD (n = 6). (C) Uninfected tissues were treated in parallel with twice daily doses of 32 J/cm^2^ of 425 nm light for three days and cytotoxicity was evaluated via MTT assay at 72 hours after the first exposure. Data presented are mean viability +/− SD (n = 6). Statistical significance was determined with the Mann-Whitney ranked sum test and is indicated by * (p≤0.05) and ** (p≤0.01).

### 425 nm light inhibits SARS-CoV-2 Delta variant infection in a human tissue model regardless of MOI

We next investigated whether the 32 J/cm^2^ BID therapeutic approach was sufficient to control SARS-CoV-2 Delta infection at multiple starting infectious titers (MOIs 0.1, 0.01, and 0.001) in the same model (Figure 6). Concordant with Beta, 32 J/cm^2^ BID reduced Delta (MOI 0.1) infectious titers by >4 log_10_ at 72 hpi; infectious Delta titers also declined from 24 hpi to 72 hpi (Figure 6A). At the lower MOIs, 425 nm light dramatically reduced infectious SARS-CoV-2 Delta after 3 days of twice daily repeat dosing (below limit of detection) (Figures 6B and 6C).

**Figure 6.**
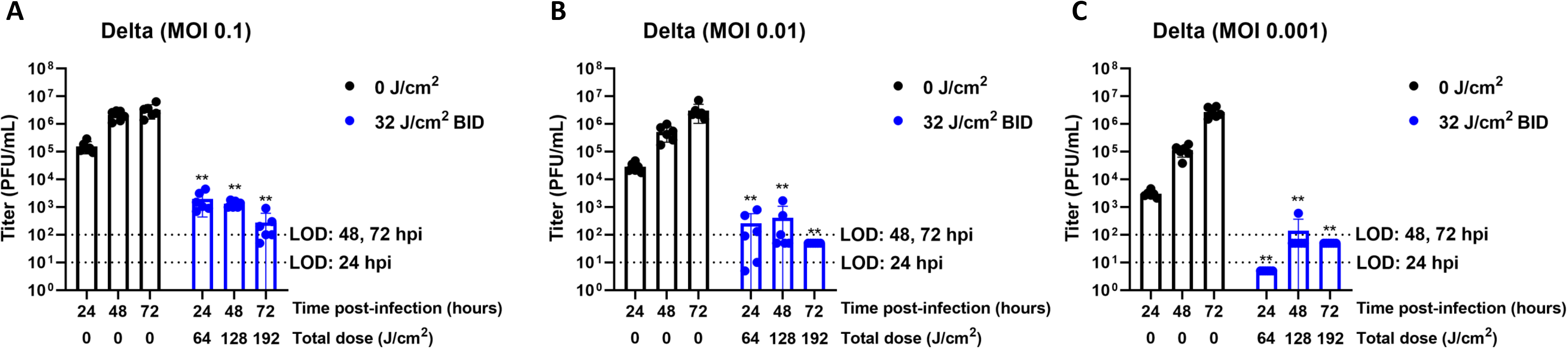
Twice daily exposures of 32 J/cm^2^ light inhibits productive SARS-CoV-2 Delta infection in the early stages of viral infection of a primary human airway tissue model. Well-differentiated primary human airway tissue models were infected with SARS-CoV-2 Delta at (A) MOI 0.1, (B) MOI 0.01, and (C) MOI 0.001 and illuminated twice daily for three days with 32 J/cm^2^ of 425 nm light starting at 3 hours post-infection. Apical rinses were collected daily and enumerated via plaque assay. Data presented are mean viral titer (PFU/mL) +/− SD (n = 6). Statistical significance was determined with the Mann-Whitney ranked sum test and is indicated by ** (p≤0.01).

The effects observed following administration of blue light during early infection (3 hpi) accurately reflect therapeutic potential early in the viral replication cycle but may not accurately reflect more established infections. To answer this question, we infected the tissue model with SARS-CoV-2 Beta or SARS-CoV-2 Delta (MOI 0.001) and delayed the first therapeutic dose to 24 hpi. Even with a delayed first dose, 32 J/cm^2^ BID effectively inhibited both variants in this model. A delayed first dose reduced Beta infectious titers by >1 log_10_ at 48 hpi, by >2 log_10_ at 72 hpi, and by >3 log_10_ at 96 hpi (Figure 7A). Similar results were seen with Delta; infectious titers were significantly reduced by >1log_10_ at 48 hpi, by >1log_10_ at 72 hpi, and by >2 log_10_ at 96 hpi (Figure 7B). These results suggest that 425 nm light therapy can inhibit SARS-CoV-2 replication at multiple stages during the viral replication cycle.

**Figure 7.**
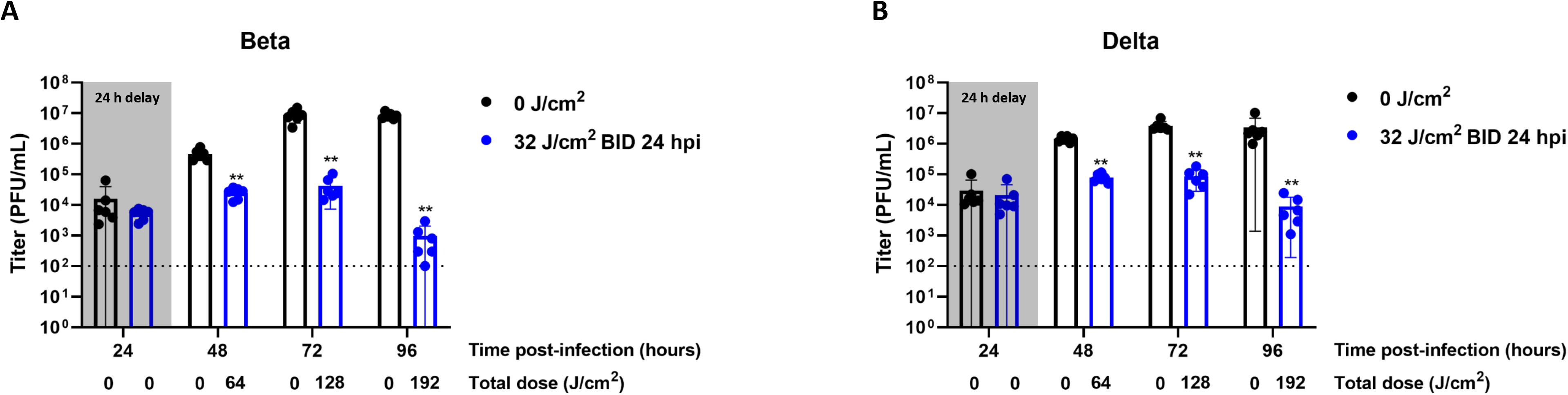
425 nm light inhibits productive SARS-CoV-2 Beta and Delta infection during later stages of viral infection in a primary human airway tissue model. Well-differentiated primary human airway tissue models were infected with SARS-CoV-2 Beta or Delta (MOI 0.001) and illuminated twice daily for 3 days with 32 J/cm^2^ of 425 nm light starting at 24 hours post-infection. Apical rinses were collected daily and enumerated via plaque assay. Data presented are mean viral titer (PFU/mL) +/− SD (n = 6) for Beta (A) and Delta (B). Statistical significance was determined with the Mann-Whitney ranked sum test and is indicated by ** (p≤0.01).

## Discussion

The rapid development and deployment of vaccines, improved standards of care, and increased focus on therapeutics have helped slow the spread of SARS-CoV-2 and the resulting worldwide economic burden. However, pockets of uncontrolled viral spread have led to the emergence of novel variants that are able to evade exiting vaccines and therapeutics (Arora et al., 2021; Cele et al., 2021; H. Liu et al., 2021; Wang et al., 2021). It is expected that more will arise. Accordingly, novel therapeutics that will work broadly against all variants, including those that have not yet emerged, are needed.

The disease state at which the novel therapeutic would be most effective must also be considered. SARS-CoV-2 infects the oral cavity, upper respiratory tract, and large airway (Hou et al., 2020; Huang et al., 2021) prior to spread to the lower respiratory tract and the late-stage development of acute respiratory distress. Sustained replication in the oral and nasal cavities is likely a key contributor to the increased transmissibility of SARS-CoV-2 compared to other coronaviruses (Hou et al., 2020; Huang et al., 2021; Marchesan et al., 2021). For these reasons, a targeted approach for acute SARS-CoV-2 infection of the upper airway epithelia to halt progression via the oral-lung transmission axis is an attractive aim. A therapeutic that works during the early stages of infection is not only essential to reduce disease burden in the treated individual, but also to limit person-to-person transmission.

Previously, we showed that 425 nm light inhibited SARS-CoV-2 WA1 in human tissue models of both oral and large airway epithelia with no damage to healthy tissue (Stasko et al., 2021b, 2021a). In this report, we show that 425 nm light can not only inactivate all SARS-CoV-2 variants of concern in cell-free suspensions, but targeted energy densities inhibit SARS-CoV-2 infections at multiple stages of infection in tissue models of human airway epithelia. Three-dimensional, differentiated primary cell culture models of human airway epithelia are more effective systems to evaluate therapeutic efficacy and safety than conventional two-dimensional immortalized cell culture systems (e.g. Vero cells) (Do et al., 2021; Heinen et al., 2021; Touret et al., 2021a). Similar models have been used in the evaluation of several anti-SARS-CoV-2 therapeutics, including molnupiravir (Sheahan et al., 2020), remdesivir (Pruijssers et al., 2020), and AT-511 (Good et al., 2021). Further, detection of infectious viruses in the apical washes can serve as a surrogate for virus shedding during infection. To this end, the proper model must be selected as different patient donors, primary cell culture conditions, and differentiation protocols can impact model development (Fulcher and Randell, 2013; Sellgren et al., 2014). For example, primary cells cultured on PTFE surfaces are more differentiated and more ciliated than primary cells cultured on PET surfaces (Sellgren et al., 2014). In our previous study, we demonstrated that the EpiAirway™ model (AIR-100) can withstand a single dose of 120 J/cm^2^ of 425 nm (Stasko et al., 2021b), but this model does not support sufficient SARS-CoV-2 WA1 replication for therapeutic exploration (Supplementary Figure 3A). The most effective model screened for these studies was developed at the Marsico Lung Institute at the University of North Carolina-Chapel Hill (Fulcher and Randell, 2013) where SARS-CoV-2 replication in the model displayed more consistent replication kinetics and higher peak titers than in the widely available AIR-100 EpiAirway™ model, providing a more robust model for therapeutic efficacy and safety evaluations in this study.

While we have previously demonstrated safety and therapeutic efficacy against WA1, several variants have replaced the parental strain. Mutations within the viral Spike, such as L18F, E484K, and N501Y, have been associated with escape from immune evasion (Cele et al., 2021; Gong et al., 2021; Harvey et al., 2021; Motozono et al., 2021; Widera et al., 2021). Herein we demonstrate inactivation of a panel of SARS-CoV-2 variants containing these mutations, indicating these mutations do not convey viral resistance to blue light inactivation. To evaluate light therapy against circulating variants of concern in a human tissue model, we selected the Beta and Delta variants due to their immune evasion and public health impacts (Pouwels et al., 2021). For both variants, early (Figures 5 and 6) or delayed (Figure 7) administration of twice daily 32 J/cm^2^ of 425 nm light for three days successfully inhibited productive infections. These results suggest that targeted 425 nm light therapy has potential treatment utility to control acute SARS-CoV-2 infections, reduce symptomatic disease and viral load, and may reduce person-to-person transmission after infection. These preclinical feasibility studies demonstrate the promising potential of 425 nm light therapy against SARS-CoV-2 variants and have led to the evaluation of an investigational treatment device currently being explored in randomized, controlled trials in outpatients with COVID-19 (“Evaluation of the RD-X19 treatment device in individuals with mild-to-moderate COVID-19.,” n.d.).

## Acknowledgments

We thank BEI Resources for providing the SARS-CoV-2 variants. We thank Dr. John McNeil, M.D. and Judy Stein for acquisition of bamlanivimab. We thank the EmitBio engineering team (Thomas M. Womble, Emily Keller, Soren Emerson, Michael Lay, P. Joseph DeSena, Haley Ritchie, and Fred Kamau) for developing and providing the biological light units. Figures 1 and 5A were created with BioRender.com

## References

Arora, P., Sidarovich, A., Krüger, N., Kempf, A., Nehlmeier, I., Graichen, L., Moldenhauer, A.-S., Winkler, M.S., Schulz, S., Jäck, H.-M., Stankov, M. v., Behrens, G.M.N., Pöhlmann, S., Hoffmann, M., 2021. B.1.617.2 enters and fuses lung cells with increased efficiency and evades antibodies induced by infection and vaccination. Cell Reports 37, 109825. https://doi.org/10.1016/j.celrep.2021.109825

Beigel, J.H., Tomashek, K.M., Dodd, L.E., Mehta, A.K., Zingman, B.S., Kalil, A.C., Hohmann, E., Chu, H.Y., Luetkemeyer, A., Kline, S., Lopez de Castilla, D., Finberg, R.W., Dierberg, K., Tapson, V., Hsieh, L., Patterson, T.F., Paredes, R., Sweeney, D.A., Short, W.R., Touloumi, G., Lye, D.C., Ohmagari, N., Oh, M., Ruiz-Palacios, G.M., Benfield, T., Fätkenheuer, G., Kortepeter, M.G., Atmar, R.L., Creech, C.B., Lundgren, J., Babiker, A.G., Pett, S., Neaton, J.D., Burgess, T.H., Bonnett, T., Green, M., Makowski, M., Osinusi, A., Nayak, S., Lane, H.C., 2020. Remdesivir for the Treatment of Covid-19 — Preliminary Report. New England Journal of Medicine. https://doi.org/10.1056/nejmoa2007764

Caly, L., Druce, J.D., Catton, M.G., Jans, D.A., Wagstaff, K.M., 2020. The FDA-approved drug ivermectin inhibits the replication of SARS-CoV-2 in vitro. Antiviral Research 178.

Cao, Y., Wang, J., Jian, F., Xiao, T., Song, W., Yisimayi, A., Huang, W., Li, Q., Wang, P., An, R., Wang, Yao, Niu, X., Yang, S., Liang, H., Sun, H., Li, T., Yu, Y., Cui, Q., Liu, S., Yang, X., Du, S., Zhang, Z., Hao, X., Shao, F., Jin, R., Wang, X., Xiao, J., Wang, Youchun, Xie, S., 2021. Omicron escapes the majority of existing SARS-CoV-2 neutralizing antibodies. Nature. https://doi.org/10.1038/s41586-021-04385-3

Cele, S., Gazy, I., Jackson, L., Hwa, S.H., Tegally, H., Lustig, G., Giandhari, J., Pillay, S., Wilkinson, E., Naidoo, Y., Karim, F., Ganga, Y., Khan, K., Bernstein, M., Balazs, A.B., Gosnell, B.I., Hanekom, W., Moosa, M.Y.S., Abrahams, S., Alcantara, L.C.J., Alisoltani-Dehkordi, A., Allam, M., Bhiman, J.N., Davies, M.A., Doolabh, D., Engelbrecht, S., Fonseca, V., Giovanetti, M., Glass, A.J., Godzik, A., Goedhals, D., Hardie, D., Hsiao, M., Iranzadeh, A., Ismail, A., Korsman, S., Pond, S.L.K., Laguda-Akingba, O., Lourenco, J., Marais, G., Martin, D., Maslo, C., Mlisana, K., Mohale, T., Msomi, N., Mudau, I., Petruccione, F., Preiser, W., San, E.J., Sewell, B.T., Tyers, L., van Zyl, G., von Gottberg, A., Walaza, S., Weaver, S., Wibmer, C.K., Williamson, C., York, D., Archary, M., Dullabh, K.J., Goulder, P., Harilall, S., Harling, G., Harrichandparsad, R., Herbst, K., Jeena, P., Khoza, T., Klein, N., Kløverpris, H., Leslie, A., Madansein, R., Marakalala, M., Mazibuko, M., Moshabela, M., Mthabela, N., Naidoo, K., Ndhlovu, Z., Ndung'u, T., Nyamande, K., Padayatchi, N., Patel, V., Ramjit, D., Rodel, H., Smit, T., Steyn, A., Wong, E., Lessells, R.J., de Oliveira, T., Sigal, A., 2021. Escape of SARS-CoV-2 501Y.V2 from neutralization by convalescent plasma. Nature 593, 142–146. https://doi.org/10.1038/s41586-021-03471-w

Cheng, Y.-W., Chao, T.-L., Li, C.-L., Wang, S.-H., Kao, H.-C., Tsai, Y.-M., Wang, H.-Y., Hsieh, C.-L., Lin, Y.-Y., Chen, P.-J., Chang, S.-Y., Yeh, S.-H., 2021. D614G Substitution of SARS-CoV-2 Spike Protein Increases Syncytium Formation and Virus Titer via Enhanced Furin-Mediated Spike Cleavage. mBio 12. https://doi.org/10.1128/mBio

Do, T.N.D., Donckers, K., Vangeel, L., Chatterjee, A.K., Gallay, P.A., Bobardt, M.D., Bilello, J.P., Cihlar, T., de Jonghe, S., Neyts, J., Jochmans, D., 2021. A robust SARS-CoV-2 replication model in primary human epithelial cells at the air liquid interface to assess antiviral agents. Antiviral Research 192. https://doi.org/10.1016/j.antiviral.2021.105122

Evaluation of the RD-X19 treatment device in individuals with mild-to-moderate COVID-19. [WWW Document], n.d.. ClinicalTrials.gov Identifier: NCT04966013. URL https://clinicaltrials.gov/ct2/show/NCT04966013 (accessed 10.31.21).

Fulcher, M.L., Randell, S.H., 2013. Human Nasal and Tracheo-Bronchial Respiratory Epithelial Cell Culture, in: Methods in Molecular Biology. pp. 109–121. https://doi.org/10.1007/978-1-62703-125-7_8

Gong, S.Y., Chatterjee, D., Richard, J., Prévost, J., Tauzin, A., Gasser, R., Bo, Y., Vézina, D., Goyette, G., Gendron-Lepage, G., Medjahed, H., Roger, M., Côté, M., Finzi, A., 2021. Contribution of single mutations to selected SARS-CoV-2 emerging variants spike antigenicity. Virology 563, 134–145. https://doi.org/10.1016/j.virol.2021.09.001

Good, S.S., Westover, J., Jung, K.H., Zhou, X.J., Moussa, A., la Colla, P., Collu, G., Canard, B., Sommadossi, J.P., 2021. AT-527, a double prodrug of a guanosine nucleotide analog, is a potent inhibitor of SARS-CoV-2 in vitro and a promising oral antiviral for treatment of covid-19. Antimicrobial Agents and Chemotherapy. 65. https://doi.org/10.1128/AAC.02479-20

Harvey, W.T., Carabelli, A.M., Jackson, B., Gupta, R.K., Thomson, E.C., Harrison, E.M., Ludden, C., Reeve, R., Rambaut, A., Peacock, S.J., Robertson, D.L., 2021. SARS-CoV-2 variants, spike mutations and immune escape. Nature Reviews Microbiology. https://doi.org/10.1038/s41579-021-00573-0

Heinen, N., Klöhn, M., Steinmann, E., Pfaender, S., 2021. In vitro lung models and their application to study sars-cov-2 pathogenesis and disease. Viruses. https://doi.org/10.3390/v13050792

Hou, Y.J., Okuda, K., Edwards, C.E., Martinez, D.R., Asakura, T., Dinnon, K.H., Kato, T., Lee, R.E., Yount, B.L., Mascenik, T.M., Chen, G., Olivier, K.N., Ghio, A., Tse, L. v., Leist, S.R., Gralinski, L.E., Schäfer, A., Dang, H., Gilmore, R., Nakano, S., Sun, L., Fulcher, M.L., Livraghi-Butrico, A., Nicely, N.I., Cameron, M., Cameron, C., Kelvin, D.J., de Silva, A., Margolis, D.M., Markmann, A., Bartelt, L., Zumwalt, R., Martinez, F.J., Salvatore, S.P., Borczuk, A., Tata, P.R., Sontake, V., Kimple, A., Jaspers, I., O'Neal, W.K., Randell, S.H., Boucher, R.C., Baric, R.S., 2020. SARS-CoV-2 Reverse Genetics Reveals a Variable Infection Gradient in the Respiratory Tract. Cell 182, 429-446.e14. https://doi.org/10.1016/j.cell.2020.05.042

Huang, N., Pérez, P., Kato, T., Mikami, Y., Okuda, K., Gilmore, R.C., Conde, C.D., Gasmi, B., Stein, S., Beach, M., Pelayo, E., Maldonado, J.O., Lafont, B.A., Jang, S.I., Nasir, N., Padilla, R.J., Murrah, V.A., Maile, R., Lovell, W., Wallet, S.M., Bowman, N.M., Meinig, S.L., Wolfgang, M.C., Choudhury, S.N., Novotny, M., Aevermann, B.D., Scheuermann, R.H., Cannon, G., Anderson, C.W., Lee, R.E., Marchesan, J.T., Bush, M., Freire, M., Kimple, A.J., Herr, D.L., Rabin, J., Grazioli, A., Das, S., French, B.N., Pranzatelli, T., Chiorini, J.A., Kleiner, D.E., Pittaluga, S., Hewitt, S.M., Burbelo, P.D., Chertow, D., Kleiner, D.E., de Melo, M.S., Dikoglu, E., Desar, S., Ylaya, K., Chung, J.Y., Smith, G., Chertow, D.S., Vannella, K.M., Ramos-Benitez, M., Ramelli, S.C., Samet, S.J., Babyak, A.L., Valenica, L.P., Richert, M.E., Hays, N., Purcell, M., Singireddy, S., Wu, J., Chung, J., Borth, A., Bowers, K., Weichold, A., Tran, D., Madathil, R.J., Krause, E.M., Herr, D.L., Rabin, J., Herrold, J.A., Tabatabai, A., Hochberg, E., Cornachione, C., Levine, A.R., McCurdy, M.T., Saharia, K.K., Chancer, Z., Mazzeffi, M.A., Richards, J.E., Eagan, J.W., Sangwan, Y., Sequeira, I., A. Teichmann, S., J. Kimple, A., Frank, K., Lee, J., Boucher, R.C., Teichmann, S.A., Warner, B.M., Byrd, K.M., 2021. SARS-CoV-2 infection of the oral cavity and saliva. Nature Medicine 27, 892–903. https://doi.org/10.1038/s41591-021-01296-8

Jayk Bernal, A., Gomes da Silva, M., Musungaie, D., Kovalchuk, E., Gonzalez, A., Delos Reyes, V., Martin-Quiros, A., Caraco, Y., Williams-Diaz, A., Brown, M., Du, J., Pedley, A., Assaid, C., Strizki, J., Grobler, J., Shamsuddin, H., Tipping, R., Wan, H., Paschke, A., Butterton, J., Johnson, M., de Anda, C, 2021. Molnupiravir for Oral Treatment of Covid-19 in Nonhospitalized Patients. The New England Journal of Medicine. https://doi.org/10.1056/NEJMoa2116044

Krause, P.R., Fleming, T.R., Longini, I.M., Peto, R., Briand, S., Heymann, D.L., Beral, V., Snape, M.D., Rees, H., Ropero, A.-M., Balicer, R.D., Cramer, J.P., Munoz-Fontela, C., Gruber, M., Gaspar, R., Singh, J.A., Subbarao, K., van Kerkhove, M.D., Swaminathan, S., Ryan, M.J., Henao-Restrepo, A.-M., 2021. SARS-CoV-2 Variants and Vaccines. The New England Journal of Medicine 385.

Kumar, S., Chandele, A., Sharma, A., 2021. Current status of therapeutic monoclonal antibodies against SARS-CoV-2 _ Enhanced Reader. PLoS Pathogens 17. https://doi.org/10.1371/journal.ppat.1009885

Levin, E.G., Lustig, Y., Cohen, C., Fluss, R., Indenbaum, V., Amit, S., Doolman, R., Asraf, K., Mendelson, E., Ziv, A., Rubin, C., Freedman, L., Kreiss, Y., Regev-Yochay, G., 2021. Waning Immune Humoral Response to BNT162b2 Covid-19 Vaccine over 6 Months. The New England Journal of Medicine.

Liu, H., Wei, P., Zhang, Q., Aviszus, K., Linderberger, J., Yang, J., Liu, J., Chen, Z., Waheed, H., Reynoso, L., Downey, G.P., Frankel, S.K., Kappler, J., Marrack, P., Zhang, G., 2021. The Lambda variant of SARS-CoV-2 has a better chance than the Delta variant to escape vaccines. bioRxiv : the preprint server for biology. https://doi.org/10.1101/2021.08.25.457692

Liu, J., Cao, R., Xu, M., Wang, X., Zhang, H., Hu, H., Li, Y., Hu, Z., Zhong, W., Wang, M., 2020. Hydroxychloroquine, a less toxic derivative of chloroquine, is effective in inhibiting SARS-CoV-2 infection in vitro. Cell Discovery. https://doi.org/10.1038/s41421-020-0156-0

Liu, Y., Liu, J., Johnson, B.A., Xia, H., Ku, Z., Widen, S.G., An, Z., Weaver, S.C., Menachery, V.D., Xie, X., Shi, P.-Y., 2021. Delta spike P681R mutation enhances SARS-CoV-2 fitness over Alpha variant 1. bioRxiv. https://doi.org/10.1101/2021.08.12.456173

Marchesan, J.T., Warner, B.M., Byrd, K.M., 2021. The “oral” history of COVID-19: Primary infection, salivary transmission, and post-acute implications. Journal of Periodontology. https://doi.org/10.1002/JPER.21-0277

McCullough, P.A., Kelly, R.J., Ruocco, G., Lerma, E., Tumlin, J., Wheelan, K.R., Katz, N., Lepor, N.E., Vijay, K., Carter, H., Singh, B., McCullough, S.P., Bhambi, B.K., Palazzuoli, A., de Ferrari, G.M., Milligan, G.P., Safder, T., Tecson, K.M., Wang, D.D., McKinnon, J.E., O'Neill, W.W., Zervos, M., Risch, H.A., 2021. Pathophysiological Basis and Rationale for Early Outpatient Treatment of SARS-CoV-2 (COVID-19) Infection. American Journal of Medicine. https://doi.org/10.1016/j.amjmed.2020.07.003

Motozono, C., Toyoda, M., Zahradnik, J., Saito, A., Nasser, H., Tan, T.S., Ngare, I., Kimura, I., Uriu, K., Kosugi, Y., Yue, Y., Shimizu, R., Ito, J., Torii, S., Yonekawa, A., Shimono, N., Nagasaki, Y., Minami, R., Toya, T., Sekiya, N., Fukuhara, T., Matsuura, Y., Schreiber, G., Ikeda, T., Nakagawa, S., Ueno, T., Sato, K., 2021. SARS-CoV-2 spike L452R variant evades cellular immunity and increases infectivity. Cell Host and Microbe 29, 1124-1136.e11. https://doi.org/10.1016/j.chom.2021.06.006

Naaber, P., Tserel, L., Kangro, K., Sepp, E., Jürjenson, V., Adamson, A., Haljasmägi, L., Rumm, A.P., Maruste, R., Kärner, J., Gerhold, J.M., Planken, A., Ustav, M., Kisand, K., Peterson, P., 2021. Dynamics of antibody response to BNT162b2 vaccine after six months: a longitudinal prospective study. The Lancet Regional Health - Europe 100208. https://doi.org/10.1016/j.lanepe.2021.100208

Planas, D., Veyer, D., Baidaliuk, A., Staropoli, I., Guivel-Benhassine, F., Rajah, M.M., Planchais, C., Porrot, F., Robillard, N., Puech, J., Prot, M., Gallais, F., Gantner, P., Velay, A., le Guen, J., Kassis-Chikhani, N., Edriss, D., Belec, L., Seve, A., Courtellemont, L., Péré, H., Hocqueloux, L., Fafi-Kremer, S., Prazuck, T., Mouquet, H., Bruel, T., Simon-Lorière, E., Rey, F.A., Schwartz, O., 2021. Reduced sensitivity of SARS-CoV-2 variant Delta to antibody neutralization. Nature 596, 276–280. https://doi.org/10.1038/s41586-021-03777-9

Plante, J.A., Liu, Y., Liu, J., Xia, H., Johnson, B.A., Lokugamage, K.G., Zhang, X., Muruato, A.E., Zou, J., Fontes-Garfias, C.R., Mirchandani, D., Scharton, D., Bilello, J.P., Ku, Z., An, Z., Kalveram, B., Freiberg, A.N., Menachery, V.D., Xie, X., Plante, K.S., Weaver, S.C., Shi, P.Y., 2021. Spike mutation D614G alters SARS-CoV-2 fitness. Nature 592, 116–121. https://doi.org/10.1038/s41586-020-2895-3

Pouwels, K.B., Pritchard, E., Matthews, P.C., Stoesser, N., Eyre, D.W., Vihta, K.-D., House, T., Hay, J., Bell, J.I., Newton, J.N., Farrar, J., Crook, D., Cook, D., Rourke, E., Studley, R., Peto, T.E.A., Diamond, I., Walker, A.S., 2021a. Effect of Delta variant on viral burden and vaccine effectiveness against new SARS-CoV-2 infections in the UK. Nature Medicine.

Pouwels, K.B., Pritchard, E., Matthews, P.C., Stoesser, N., Eyre, D.W., Vihta, K.-D., House, T., Hay, J., Bell, J.I., Newton, J.N., Farrar, J., Crook, D., Cook, D., Rourke, E., Studley, R., Peto, T.E.A., Diamond, I., Walker, A.S., 2021b. Effect of Delta variant on viral burden and vaccine effectiveness against new SARS-CoV-2 infections in the UK. Nature Medicine. https://doi.org/10.1038/s41591-021-01548-7

Pruijssers, A.J., George, A.S., Schäfer, A., Leist, S.R., Gralinksi, L.E., Dinnon, K.H., Yount, B.L., Agostini, M.L., Stevens, L.J., Chappell, J.D., Lu, X., Hughes, T.M., Gully, K., Martinez, D.R., Brown, A.J., Graham, R.L., Perry, J.K., du Pont, V., Pitts, J., Ma, B., Babusis, D., Murakami, E., Feng, J.Y., Bilello, J.P., Porter, D.P., Cihlar, T., Baric, R.S., Denison, M.R., Sheahan, T.P., 2020. Remdesivir Inhibits SARS-CoV-2 in Human Lung Cells and Chimeric SARS-CoV Expressing the SARS-CoV-2 RNA Polymerase in Mice. Cell Reports 32, 107940. https://doi.org/10.1016/j.celrep.2020.107940

Ren, S.-Y., Wang, W.-B., Gao, R.-D., Zhou, A.-M., 2022. World Journal of Clinical Cases Omicron variant (B.1.1.529) of SARS-CoV-2: Mutation, infectivity, transmission, and vaccine resistance. World J Clin Cases 10, 1–11. https://doi.org/10.12998/wjcc.v10.i1.1

Ritchie, H., Mathieu, E., Rodes-Guirao, L., Appel, C., Giattino, C., Ortiz-Ospina, E., Hasell, J., Macdonald, B., Beltekian, D., Roser, M., 2020. Coronavirus Pandemic (COVID-19) [WWW Document]. OurWorldinData.org.

Sellgren, K.L., Butala, E.J., Gilmour, B.P., Randell, S.H., Grego, S., 2014. A biomimetic multicellular model of the airways using primary human cells. Lab Chip 14. https://doi.org/10.1039/C4LC00552J

Sheahan, T.P., Sims, A.C., Zhou, S., Graham, R.L., Pruijssers, A.J., Agostini, M.L., Leist, S.R., Schäfer, A., Dinnon Iii, K.H., Stevens, L.J., Chappell, J.D., Lu, X., Hughes, T.M., George, A.S., Hill, C.S., Montgomery, S.A., Brown, A.J., Bluemling, G.R., Natchus, M.G., Saindane, M., Kolykhalov, A.A., Painter, G., Harcourt, J., Tamin, A., Thornburg, N.J., Swanstrom, R., Denison, M.R., Baric, R.S., 2020. An orally bioavailable broad-spectrum antiviral inhibits SARS-CoV-2 in human airway epithelial cell cultures and multiple coronaviruses in mice, Sci. Transl. Med.

Stasko, N., Cockrell, A.S., Kocher, J.F., Henson, I., Emerson, D., Wang, Y., Smith, J.R., Henderson, N.H., Wood, H., Bradrick, S.S., Jones, T.M., Santander, J.L., McNeil, J.G., 2021a. A randomized, controlled, feasibility study of RD-X19 in patients with mild-to-moderate COVID-19 in the outpatient setting. medRxiv: the preprint server for health sciences.

Stasko, N., Kocher, J.F., Annas, A., Henson, I., Seitz, T.S., Miller, J., Arwood, L., Roberts, R.C., Womble, T.M., Keller, E.G., Emerson, S., Bergmann, M., Sheesley, A.N.Y., Strong, R.J., Hurst, B.L., Emerson, D., Tarbet, E.B., Bradrick, S.S., Cockrell, A.S., 2021b. Visible blue light inhibits infection and replication of SARS-CoV-2 at doses that are well-tolerated by human respiratory tissue. Scientific Reports 11.

Touret, F., Driouich, J.S., Cochin, M., Petit, P.R., Gilles, M., Barthélémy, K., Moureau, G., Mahon, F.X., Malvy, D., Solas, C., de Lamballerie, X., Nougairède, A., 2021a. Preclinical evaluation of Imatinib does not support its use as an antiviral drug against SARS-CoV-2. Antiviral Research 193. https://doi.org/10.1016/j.antiviral.2021.105137

Touret, F., Luciani, L., Baronti, C., Cochin, M., Driouich, J.S., Gilles, M., Thirion, L., Nougairède, A., de Lamballerie, X., 2021b. Replicative fitness of a sars-cov-2 20i/501y.V1 variant from lineage b.1.1.7 in human reconstituted bronchial epithelium. mBio 12. https://doi.org/10.1128/mBio.00850-21

VanBlargan, L.A., Errico, J.M., Halfmann, P.J., Zost, S.J., Crowe, J.E., Purcell, L.A., Kawaoka, Y., Corti, D., Fremont, D.H., Diamond, M.S., 2022. An infectious SARS-CoV-2 B.1.1.529 Omicron virus escapes neutralization by therapeutic monoclonal antibodies. Nature Medicine 1–6. https://doi.org/10.1038/s41591-021-01678-y

Wang, P., Nair, M.S., Liu, L., Iketani, S., Luo, Y., Guo, Y., Wang, M., Yu, J., Zhang, B., Kwong, P.D., Graham, B.S., Mascola, J.R., Chang, J.Y., Yin, M.T., Sobieszczyk, M., Kyratsous, C.A., Shapiro, L., Sheng, Z., Huang, Y., Ho, D.D., 2021. Antibody resistance of SARS-CoV-2 variants B.1.351 and B.1.1.7. Nature 593, 130–135. https://doi.org/10.1038/s41586-021-03398-2

Widera, M., Wilhelm, A., Hoehl, S., Pallas, C., Kohmer, N., Wolf, T., Rabenau, H.F., Corman, V.M., Drosten, C., Vehreschild, M.J.G.T., Goetsch, U., Gottschalk, R., Ciesek, S., 2021. Limited Neutralization of Authentic Severe Acute Respiratory Syndrome Coronavirus 2 Variants Carrying E484K In Vitro. The Journal of Infectious Diseases. https://doi.org/10.1093/infdis/jiab355

Xu, Z., Liu, K., Gao, G.F., 2021. Omicron variant of SARS-CoV-2 imposes a new challenge for the global pub-lic health. https://doi.org/10.1016/j.bsheal.2022.01.002

