## Supplementary Figure Legends for "Visible blue light inactivates SARS-CoV-2 variants and inhibits Delta replication in differentiated human airway epithelia"

**Supplementary Figure 1. Basal media formulation does not impact SARS-CoV-2 Beta inactivation by 425 nm light.** SARS-CoV-2 Beta was propagated in basal media composed of (A) MEM and (B) DMEM with different serum supplementation. Cell-free suspensions of SARS-CoV-2 variant Beta (n = 4) were illuminated with 425 nm light and enumerated via plaque assay. Data presented are mean viral titers (PFU/mL) +/- SD (n = 4). Statistical significance was determined via Mann-Whitney ranked sum test and is indicated by * ($p\leq 0.05$).

**Supplementary Figure 2. 425 nm light inhibits cell-free SARS-CoV-2 cell entry in an ACE-2-dependent manner.** Vero E6 cells were inoculated with cell-free suspensions of SARS-CoV-2 Beta following illumination with 0 J/cm$^2$, 15 J/cm$^2$, and 90 J/cm$^2$ of 425 nm light. At 3 hpi and 24 hpi, total RNA was extracted from inoculated cultures for qRT-PCR analysis of N2 and RPP (n = 4). Data presented are mean Ct +/- SD (n = 4) for the (A) N2 probe at 3 hpi and (B) 24 hpi and (C) normalized N1 expression relative to RNase P and the RNase P probe at (D) 3 hpi and (E) 24 hpi. Statistical significance was determined via the Mann-Whitney ranked sum test and is indicated by * ($p\leq 0.05$) and ** ($p\leq 0.01$).

**Supplementary Figure 3. SARS-CoV-2 WA1 replication in different tracheobronchial tissue models and susceptibility to 425 nm blue light.** Tracheobronchial epithelial models (AIR-100, TBE-14 and DD065Q, n = 4-6) were infected with SARS-CoV-2 WA1 (MOIs 1, 0.5 and 0.2, respectively). Cultures with productive infection were illuminated once daily with 0 J/cm$^2$, 16 J/cm$^2$, and 32 J/cm$^2$ of 425 nm light starting at 3 hpi. Apical rinses were collected at 24 hpi, 48 hpi, and 72 hpi and enumerated via plaque assay. Data presented are mean viral titers (PFU/mL) at each timepoint +/- SD for AIR-100 (A), TBE-14 (B), and DD065Q (C) tissues. Statistical significance was determined via the Mann-Whitney ranked sum test and is indicated by * ($p\leq 0.05$) and ** ($p\leq 0.01$).
