## Supplementary Figures for "Visible blue light inactivates SARS-CoV-2 variants and inhibits Delta replication in differentiated human airway epithelia"

**Supplementary Figure 1. Basal media formulation does not impact SARS-CoV-2 Beta inactivation by 425 nm light.**

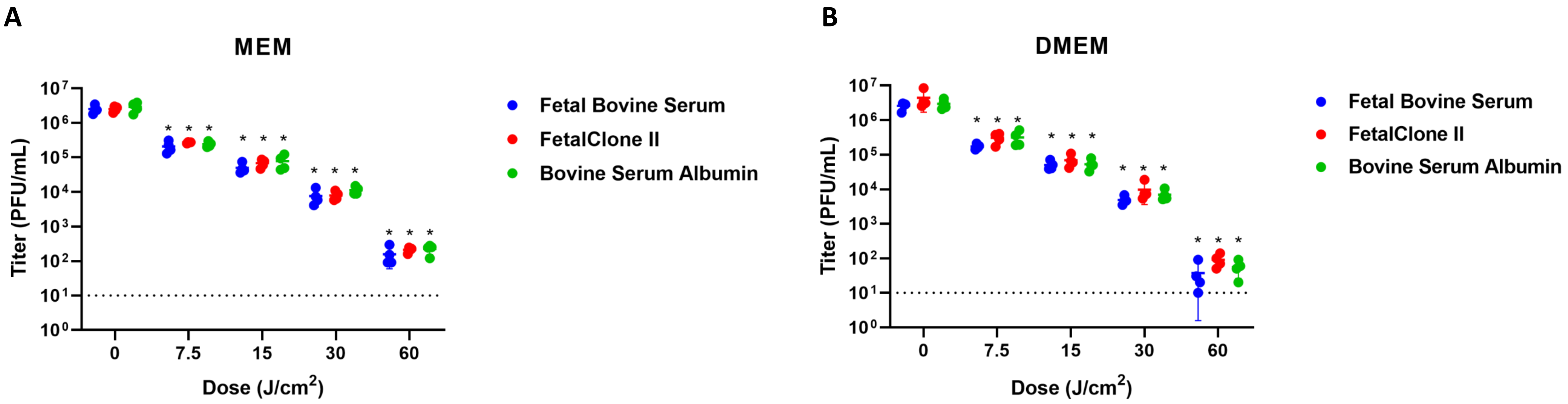

**Supplementary Figure 2. 425 nm light inhibits cell-free SARS-CoV-2 cell entry in an ACE-2-dependent manner.**

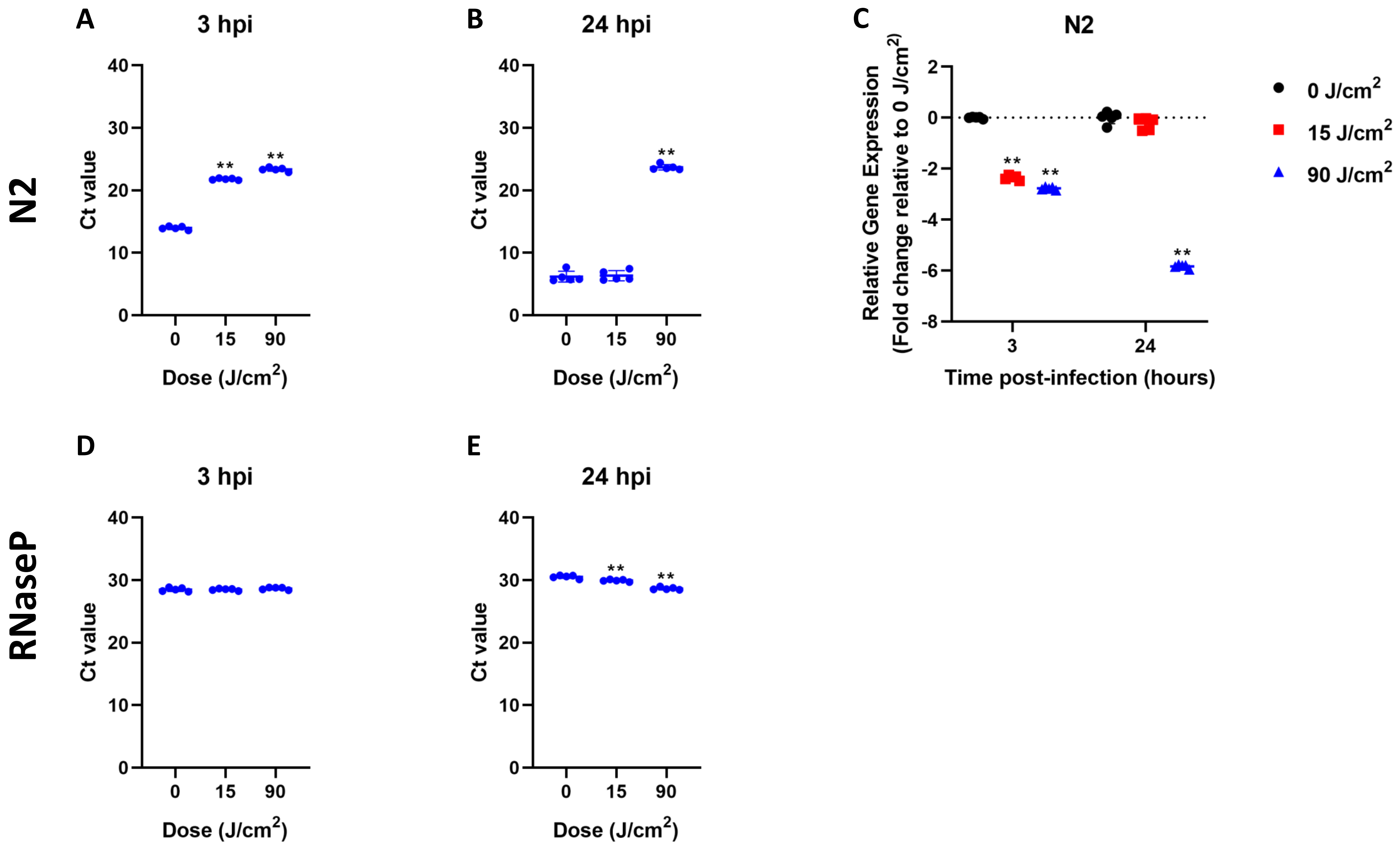

**Supplementary Figure 3. SARS-CoV-2 WA1 replication in different tracheobronchial tissue models and susceptibility to 425 nm blue light.**

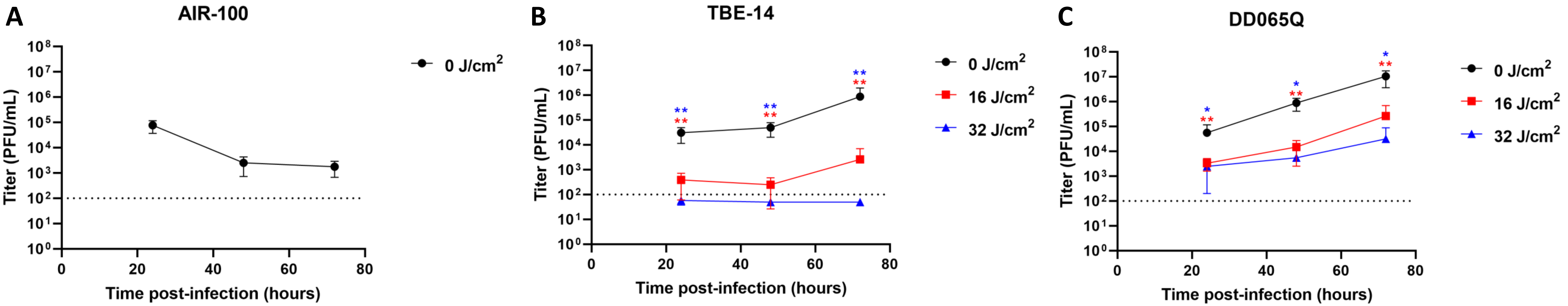
